# Genetic and selective constraints on the optimization of gene product diversity

**DOI:** 10.1101/2024.07.17.603951

**Authors:** Daohan Jiang, Nevraj Kejiou, Yi Qiu, Alexander F. Palazzo, Matt Pennell

## Abstract

RNA and protein expressed from the same gene can have diverse isoforms due to various post-transcriptional and post-translational modifications. For the vast majority of alternative isoforms, It is unknown whether they are adaptive or simply biological noise. As we cannot experimentally probe the function of each isoform, we can ask whether the distribution of isoforms across genes and across species is consistent with expectations from different evolutionary processes. However, there is currently no theoretical framework that can generate such predictions. To address this, we developed a mathematical model where isoform abundances are determined collectively by *cis*-acting loci, *trans*-acting factors, gene expression levels, and isoform decay rates to predict isoform abundance distributions across species and genes in the face of mutation, genetic drift, and selection. We found that factors beyond selection, such as effective population size and the number of *cis*-acting loci, significantly influence evolutionary outcomes. Notably, suboptimal phenotypes are more likely to evolve when the population is small and/or when the number of *cis*-loci is large. We also explored scenarios where modification processes have both beneficial and detrimental effects, revealing a non-monotonic relationship between effective population size and optimization, demonstrating how opposing selection pressures on *cis*- and *trans*-acting loci can constrain the optimization of gene product diversity. As a demonstration of the power of our theory, we compared the expected distribution of A-to-I RNA editing levels in coleoids and found this to be largely consistent with non-adaptive explanations.

## Introduction

Different RNA and protein isoforms can be expressed from the same gene, resulting in a phenomenon known as gene product diversity [1]. A variety of processes can generate gene product diversity, such as alternative transcription initiation [2, 3], alternative splicing [4–8], alternative polyadenylation [9], post-transcriptional RNA modifications [10–13], alternative translation initiation [14], post-translational modifications [8, 15], and errors during RNA or protein synthesis [16–19]. The growing body of transcriptomic and proteomic data has unveiled substantial gene product diversity produced by different processes in diverse taxa, but functional significance of the alternative isoforms remains largely unknown [1, 7, 8, 12, 13].

One explanation for observed gene product diversity is the adaptive hypothesis that the alternative isoforms perform essential functions and are beneficial to the organism [19–21]. Cases of beneficial gene product modifications have been documented in various taxa. Notable examples of potentially adaptive modification events include a nonsynonymous A-to-I RNA editing event in a potassium channel protein that confers cold tolerance in polar octopuses [22], A-to-I editing events in filamentous fungi that fix premature stop codons in proteins involved in sexual reproduction [23, 24], alternative splicing of *Sxl* transcripts that regulate sex determination in dipteran insects [25], and some circular RNA isoforms that function as micro RNA in sponges [26, 27]. However, such cases collectively comprise only a small portion of known gene product diversity.

An alternative view suggests that gene product diversity is largely non-adaptive and reflect errors in biochemical processes. Gene product modification processes that result in gene product diversity, like all other biochemical reactions, are fundamentally stochastic and thus prone to errors. While natural selection can act to reduce the error rate, optimization will be limited by a drift barrier: at small effective population sizes, molecular errors with mild fitness effect cannot be purged efficiently by selection in the face of strong genetic drift and/or mutational pressure [28, 29]. This view has been supported by analyses of various types of gene product diversity, such as alternative splicing [30–33], alternative polyadenylation [34], A-to-I RNA editing [35–37], and C-to-U RNA editing [38]. It is also plausible that different isoforms of a gene’s product are functionally equivalent, in which case the diversity *per se* is not adaptive even if the process that generates diversity is. That is, it is the amount of modification in a molecule rather than the precise location of any modification that matters. Processes that can potentially generate such neutral diversity include N6-methyladenosine (m6A) modification of RNA [38–40] and protein phosphorylation [41, 42].

Furthermore, a machinery that generates gene product diversity can be maintained by making other-wise strongly deleterious mutations reasonably benign. In such a case, it could become indispensable over time as more such mutations are permitted and fixed, a process, known as entrenchment or “constructive neutral evolution” [43–46]. For example, A-to-I editing can “permit” G-to-A mutations as inosine (I) is recognized as guanine (G) during translation; this harm-permitting effect has likely contributed to maintenance of high A-to-I editing activity in coleoid cephalopods (subclass Coleoidea, including octopuses, squids, and cuttlefishes) [36]. Similarly, high C-to-U editing in plant organelles may have been entrenched after permitting T-to-C mutations [47–49].

One possible way to distinguish these alternative hypotheses in the absence of functional information for the vast majority of isoforms is to compare the observed gene product diversity within and between species to that expected under various evolutionary scenarios. However, such comparisons are not currently possible as we lack a theoretical basis for generating such expectations. While phylogenetic comparative methods have recently been applied to molecular phenotypes like gene expression levels (e.g., [50–54]), it is unclear whether conventional trait evolution models used in phylogenetic comparative analyses are suitable for modeling gene product diversity. To address these, we developed a mathematical model that connects patterns of variation in gene product diversity and the underlying evolutionary processes. In particular, we investigated two types of gene product modification processes that represent a broad range of processes that generate gene product diversity. The first type of modification simply converts an unmodified isoform to modified isoform(s) that can potentially be dysfunctional and/or toxic (Fig. 1A). Such modifications are not universally required for gene products to carry out their primary functions. Prime examples of such modifications include a variety of post-transcriptional RNA editing processes, where the RNA molecule is enzymatically modified into an alternative isoform [10–13]. Thus, we will refer to this type of processes as “editing-type”. The second type of gene product modification process is required to produce the functional isoform, but can potentially produce mis-processed isoforms that could be dysfunctional and/or toxic (Fig. 1B). This class of modification is exemplified by RNA splicing in eukaryotes, which is generally required but can potentially produce toxic mis-spliced isoforms [4, 5]. Thus, this second type of gene product modification is referred to as “splicing-type”. In both cases, each gene product modification event is regulated by a set of *cis*-loci and a *trans*-factor. Each *cis*-locus only affects a specific modification event and thus has a local effect, whereas the *trans*-factor globally affects many modification events.

**Figure 1:**
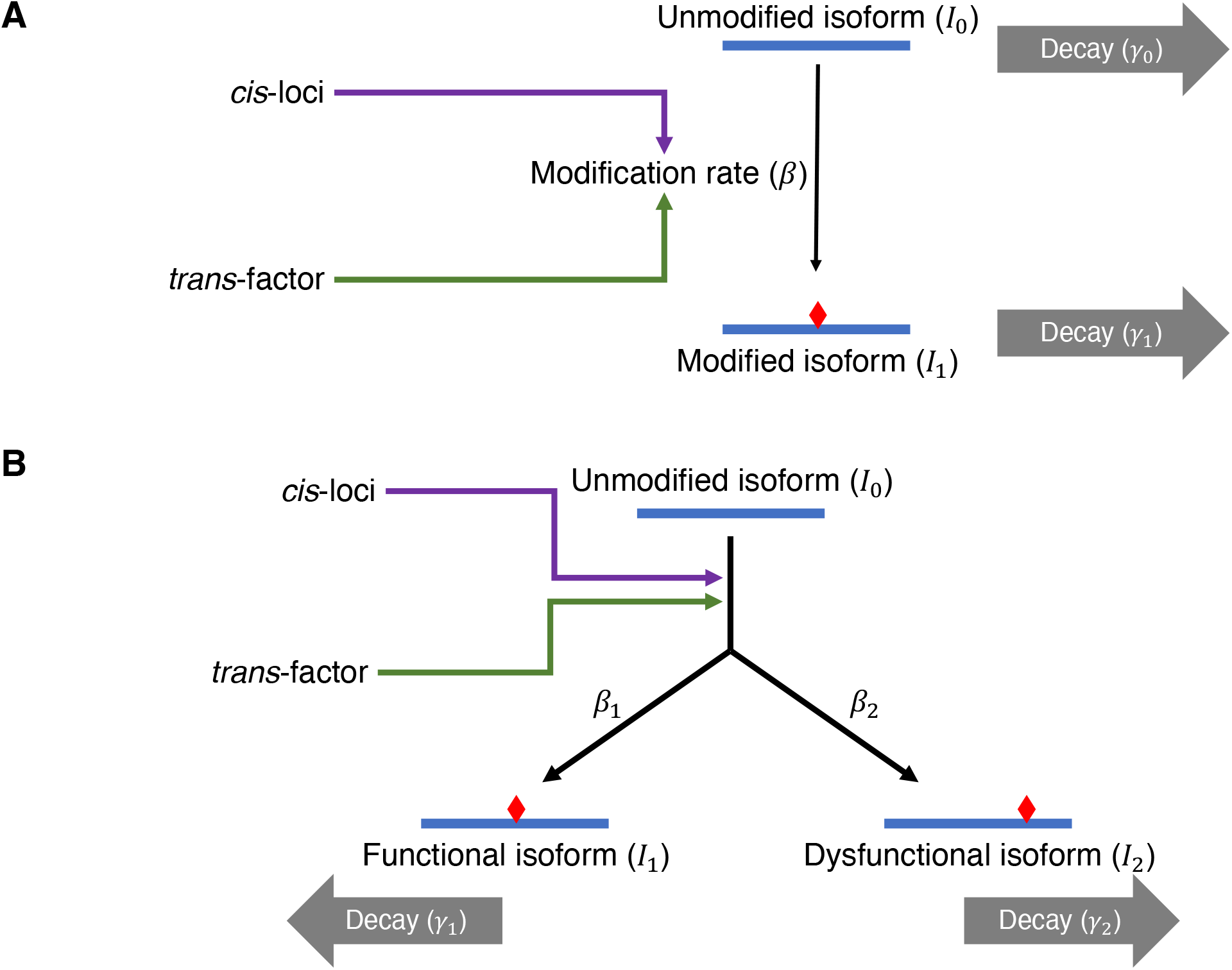
A schematic illustration of editing-type (A) and splicing-type (B) gene product diversity. (A) An unmodified isoform (*I*_0_) is enzymatically converted to a modified isoform (*I*_1_). The net per-molecule conversion rate (*β*) is determined jointly by a *trans*-factor (enzyme performing the modification process) and a set of *cis*-loci (sequence motif underlying affinity between enzyme and substrate). (B) The unmodified isoform *I*_0_ can be converted into either a functional isoform (*I*_1_) or a dysfunctional isoform (*I*_2_) through the same modification process such that two conversion rates *β*_1_ and *β*_2_ are affected by the same *cis*-loci and *trans*-factor.

Under our model, we derived phylogenetic means of the modification level (i.e., relative abundance of modified isoforms) under different conditions, demonstrating how modification level is shaped by mutational pressure, genetic drift, and selection. We also investigated how opposing selection on the modification process shapes the coevolution of *cis*- and *trans*-acting loci underlying modification. At last, using computer simulations, we demonstrated that our model can recapitulate distribution of A-to-I RNA editing levels observed in empirical studies.

## Results and Discussion

### Modeling genetic architecture of isoform abundances

Under a simple model where an unmodified isoform, *I*_0_, is converted to a modified isoform, *I*_1_, rates at which their abundances in the cell changes over time can be written as

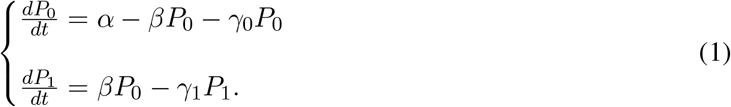

Here, *P*_0_ and *P*_1_ are abundances of *I*_0_ and *I*_1_, respectively, *α* is the rate at which *I*_0_ is produced, *β* is the per-molecule net rate at which *I*_0_ is converted to *I*_1_, and *γ*_0_ and *γ*_1_ are *I*_0_ and *I*_1_’s respective decay rates. An equilibrium is reached when both rates are equal to zero:

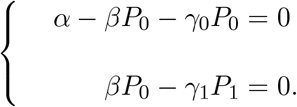

Solving the system of equations gives equilibrium isoform abundances:

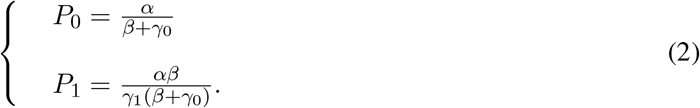

The same modeling approach can be generally applied to systems with more isoforms—e.g., *I*_0_ is converted to more than one modified isoforms (see Methods).

In our model, the per-molecule conversion rate *β* is controlled by a *trans*-factor (i.e., enzyme that performs gene product modification) and a set of *cis*-loci (i.e., sequence motif modulating enzyme binding affinity). The *trans*-factor’s effect on *β* is characterized by a *trans*-genotypic value, *Q*, which reflects the modification enzyme’s expression level and/or catalysis efficiency. The *cis*-genotype’s effect is summarized by a normalized *cis*-genotypic value 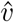. A high 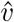 indicates strong binding between the modification enzyme and the substrate, which results in high modification efficiency, whereas a low 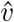 means weak enzyme-substrate binding and low modification efficiency. Each *cis*-locus can have either an effector allele that facilitates enzyme binding, or a null allele that has no effect. In this study, we focused on a simple model where all loci’s effector alleles have an equal, additive effect [29], so 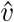 is calculated as 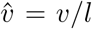, where *l* is the number of *cis*-loci that affect the modification and *v* is the total number of effector alleles. While this study is focused on such a simple model, it can readily be extended to incorporate variation in different loci’s contribution–for example, a skewed distribution where one locus has major effect while others’ effects are much weaker.

Given values of *Q* and 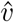, *β* is calculated as

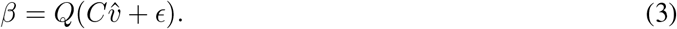

Here, *C >* 0 represents whole-molecule features that modulate the *cis*-loci’s effect size (e.g., secondary structure of RNA or protein), and *ϵ* ≥ 0 is the rate of non-specific modification (i.e., promiscuous activity of the enzyme independent of the *cis*-genotype).

For editing-type modification, we focused on a simple scenario where two isoforms, the unmodified isoform *I*_0_ and modified isoform *I*_1_, are present (Fig. 1A); the generic, two-isoform model described above is thus readily applicable. We considered values of *l* that are relatively small (i.e., no more than 10), as empirical studies suggest that sequence motifs with major effects on RNA modifications usually consist of a small number of nucleotide sites [10, 13, 55, 56]. In an extreme case, A-to-I editing in filamentous fungi, the nucleotide site immediately upstream the editable A site appears to be the only *cis*-locus, where the effector allele is a T base [23, 57, 58].

For splicing-type modification, we considered a model where the unmodified isoform *I*_0_ is converted to two modified isoforms, a functional isoform *I*_1_ and a dysfunctional isoform *I*_2_, at rates *β*_1_ and *β*_2_, respectively. As *I*_1_ and *I*_2_ are essentially products of the same process, their respective modification rates *β*_1_ and *β*_2_ are controlled by the same *cis*-loci (Fig. 1B); thus, we assumed an allele that does not facilitate production of the *I*_1_ will facilitate production of *I*_2_ and vice versa. For convenience, the *cis*-genotypic value is defined as the *cis*-genotype’s effect on *β*_1_ for splicing-type modification. Hence, there is

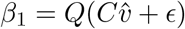

and

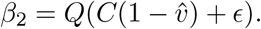

As splicing of a gene’s transcript can be affected by a relatively large number of loci, including splicing enhancers, inhibitors, and cryptic splice sites [59, 60], we considered relatively large values of *l* (10, 20, 30, 40, and 50) for splicing-type modification. We assumed *γ*_0_ = 0 but a high *Q* such that the *I*_0_ only comprise a small fraction of of gene product (i.e., *P*_0_*/*(*P*_0_ + *P*_1_ + *P*_2_) ≈ 1%) to recapitulate the fact that splicing takes place co-transcriptionally [61]. We also had *γ*_2_ significantly greater than *γ*_1_ to reflect the effect of quality control processes, such as nonsense-mediated RNA decay [62–64], or nuclear retention and decay of intronic polyadenylated transcripts mediated by recognition of intact 5’-splice site [65–67]. The model for splicing-type modification can be readily applied as long as gene product diversity results from alternative products of an indispensable process in gene expression. For instance, it may be applied to alternative polyadenylation, in which case *I*_0_ represents nascent RNA, and *I*_1_ and *I*_2_ represent RNAs polyadenylated at different sites.

### Evolutionary scaling of mean modification level

When the only loci that evolve are the *cis*-loci (e.g., the *trans*-factor is invariable because of its pleiotropic effects) and the *cis*-loci’s fitness effect is only mediated by gene product modification (i.e., the *cis*-loci have no pleiotropic effects), evolution of the *cis*-genotypic value *v* can be modeled as a discrete-state Markov process, and we can derive the probability distribution of *v* (and 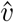) given the initial distribution and regime of selection after evolution for a given amount of time [28, 29]. To this end, we asked what the expected relative abundance of a dysfunctional, toxic isoform (e.g., reduce fitness due to mis-interactions with other biomolecules) will be in the face of mutation, drift, and selection.

For editing-type modification, we considered a deleterious modification event that converts an unmodified isoform *I*_0_ that is functional (i.e., *P*_0_ under stabilizing selection and fitness is a Gaussian function of ln *P*_0_; see Methods) to a modified isoform *I*_1_ that is not functional but toxic (i.e., fitness declines with *P*_1_; see Methods). Specifically, we examined the modification level, *f* = *P*_1_*/*(*P*_0_ + *P*_1_). For each combination of parameter values, we calculated the mean of *v* after evolution from *v* = 0 for 10^8^ time steps (i.e., generations) and the corresponding *f*, which we refer to as a phylogenetic mean of modification level (mean modification level, for short). Under all conditions examined, the mean modification level declines with effective population size *N*_*e*_ (Fig. 2). Mutational bias towards the effector allele makes the mean modification level higher, whereas bias in an opposite direction makes it lower (Fig. 2A-C). For a given *N*_*e*_ and the per-locus mutation rate, the mean modification level becomes higher when the number of *cis*-loci, *l*, is high, which is most pronounced at relatively small *N*_*e*_ (Fig. 2A-C). This relationship between modification *l* is explained by the relative size of genotypic space that produce the optimal phenotype. The optimal genotype, which leads to *v* = 0, corresponds to 2^−*l*^ of the genotypic space. Thus, when *l* is large, it is harder to maintain an optimal genotype in the face of mutational pressure towards non-zero *cis*-genotypic values when *l* is greater [29].

**Figure 2:**
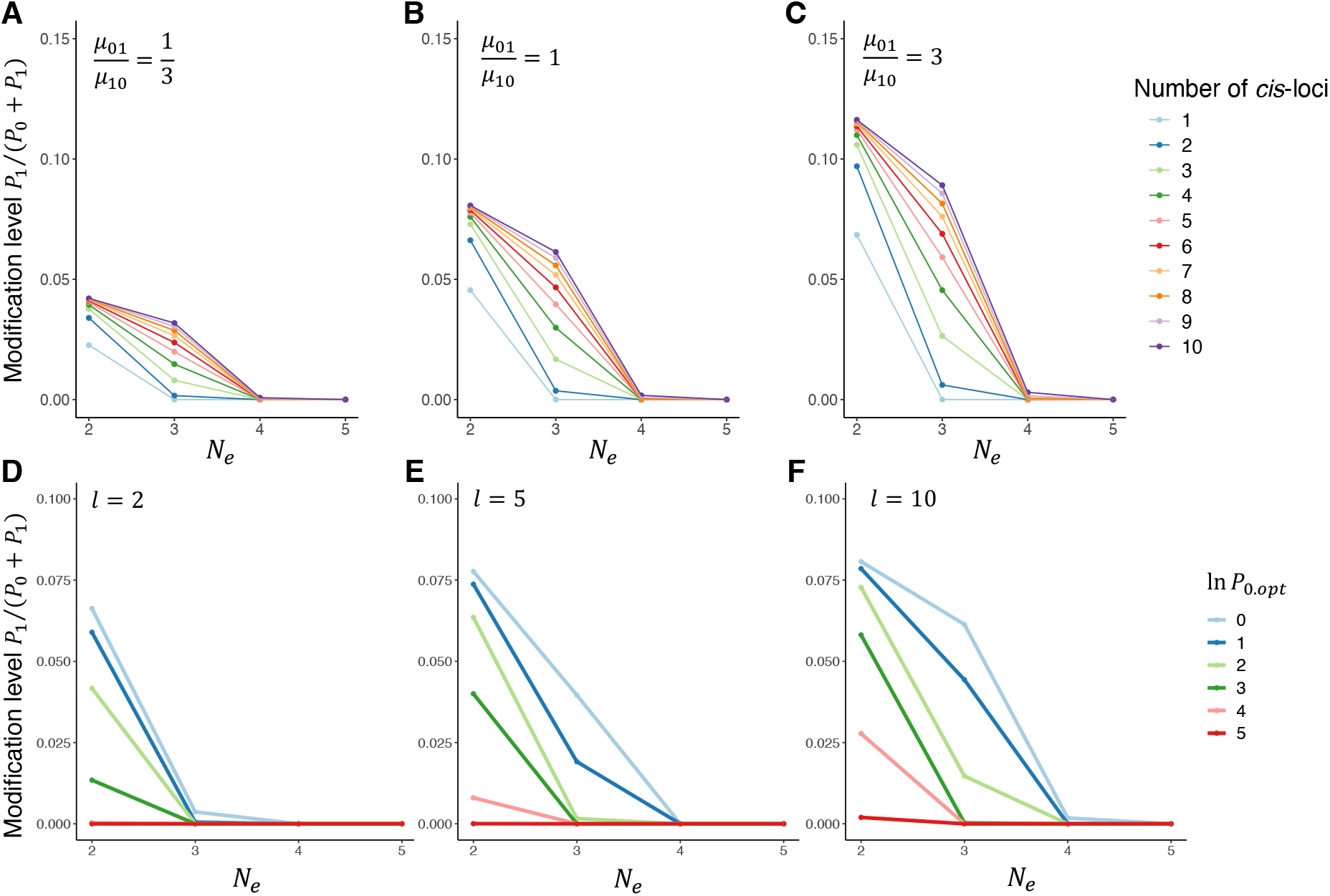
Scaling between mean modification level of a deleterious editing-type modification to effective population size *N*_*e*_ (shown in log10 scale). (A-C) Response of mean modification level to *N*_*e*_ given different combinations of *cis*-loci number (*l*) and mutation rates (*μ*_01_, *μ*_10_), with optimal expression level *P*_0.*opt*_ = exp (1) (ln *P*_0.*opt*_ = 1). (A) Mutational bias is towards the null allele that does not facilitate modification. (B) Mutations of two directions have equal mutation rates. (C) Mutational bias is towards the effector allele that facilitates modification. (D-F) Response of mean modification level to *N*_*e*_ given different *P*_0.*opt*_ with *l* = 2 (D), *l* = 5 (E), and *l* = 10 (F) in the absence of mutational bias. All results are derived with initial *cis*-genotypic value *v*_0_ = 0, time of evolution *T* = 10^8^ time steps, total mutation rate per *cis*-locus *μ* = *μ*_01_ + *μ*_10_ = 2 *×* 10^−9^, *Q* = 1, *γ*_0_ = 1, and *γ*_1_ = 1. Optimal expression level *P*_0.*opt*_ is set to be equal to *P*_0_ in the absence of modification (i.e.,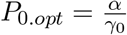) in all cases.

Another key factor affecting the mean modification level is expression level of the gene (i.e., optimal *P*_0_, reached when *β* = 0): mean modification level is lower when the gene is more highly expressed (Fig. 2D-F). This relationship is explained by toxic effect of *I*_1_–given the modification level, there will be higher *P*_1_ and thus greater fitness cost mediated by toxicity when the gene is highly expressed. It is a general phenomenon that high expression levels magnify the impact of errors in gene products and lead to in stronger selective constraints, as supported by previous studies of sequence evolution [68, 69], expression level [70], translation fidelity [71], as well as several types of gene product modifications [32–34].

For splicing-type modification, modification level is defined as relative abundance of the dysfunctional and toxic isoform *I*_2_ out of all modified products, *f* = *P*_2_*/*(*P*_1_ + *P*_2_). As in the case of editing-type modification, mean level of splicing-type modification also declined with *N*_*e*_ and gene expression level, and increased with *l* (Fig. S1). We also compared the effect of a quality-control mechanism like nonsense-mediated decay (i.e., high *γ*_2_) and confirmed that faster decay of *I*_2_ can substantially lower the modification (Fig. S1). When it is the *cis*-genotypic value that is examined, results under different values of *γ*_2_ are mostly similar (Fig. S2); when the gene’s expression level is high (i.e., optimal *P*_1_ is exp (4) or exp (5)) and *N*_*e*_ is intermediate (i.e., 10^3^-10^4^), high *γ*_2_ will have a harm-permitting effect: *cis*-genotypes that lead to production of more *I*_2_ will be permitted as the harmful effect is reduced by fast decay of *I*_2_ (Fig. S2D-F, red and pink curves).

### Non-monotonic scaling in *cis*-*trans* coevolution

Given that non-adaptive gene product diversity will be present when selection is unable to optimize the *cis*-loci in the face of mutational pressure and genetic drift, obvious questions are: why did this machinery evolve in the first place?; and how is this maintained? These are particular pertinent for editing-type modification that is not an indispensable part of gene expression. Presumably, such gene product modification processes must have additional essential functions unrelated to the set of modification events studied here, such that loss or suppression of the modification machinery will have a strongly deleterious effect. To better understand evolutionary dynamics when the modification machinery is under opposing selection forces, we considered a scenario where modification events under concern are deleterious but the *trans*-genotypic value *Q* is under stabilizing selection due to its contribution to an additional fitness component (Fig. 3A; also see Methods), and conducted simulations to investigate how *cis*- and *trans*-acting loci will respond to selection. We simulated evolution under different combinations of *N*_*e*_, *l*, and strength of selection on *Q*. The simulation started from a high value of *Q* and intermediate *cis*-genotypic values (i.e., values with the largest corresponding genotypic space), representing a state that high modification activity had just evolved and optimization of *cis*-loci have not yet started.

**Figure 3:**
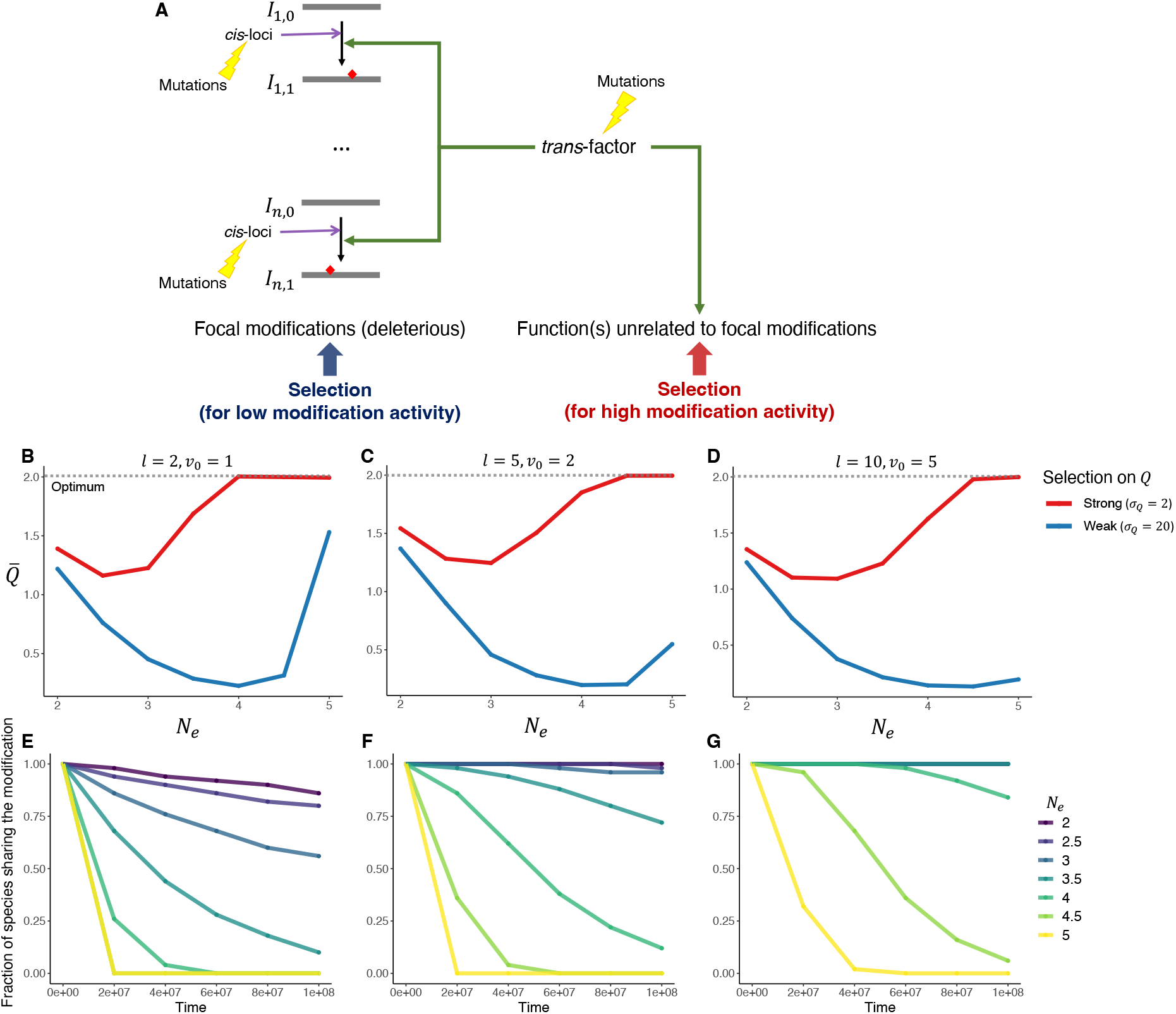
Coevolution of *cis*- and *trans*-acting loci when the gene product modification machinery is under opposing selection forces. (A) Schematic illustration of the scenario. The *trans*-factor, while causing a number of deleterious editing-type modification events (focal modifications), also performs an essential function independent of the focal modifications. Selection against deleterious modification may act to reduce the *trans*-genotypic value (*Q*), while selection mediated by the other function(s) act to maintain an optimal value of *Q* (*Q*_*opt*_). (B-D) Non-monotonic response of mean of *Q* across lineages to *N*_*e*_ (shown in log10 scale) with *Q* under stabilizing selection and 100 genes subject to deleterious modification. Curves of different colors correspond to scenarios of strong (red) and weak (blue) selection on *Q*. Optimum of *Q* is denoted by the dashed line. All simulations started with an intermediate *cis*-genotypic value with largest corresponding genotypic space. (E-G) Sharing of modification events over time. Y-axes represent among-gene median of proportion of lineages (species) that share a modification event when selection on *Q* is strong (*σ*_*Q*_ = 2). When two curves in the same panel completely overlap, the one with the largest corresponding *N*_*e*_ is shown. In (B) and (E), *l* = 2 and *v*_0_ = 1; in (C) and (F), *l* = 5 and *v*_0_ = 2; in (D) and (G), *l* = 10 and *v*_0_ = 5.

We found the among-lineage average of *Q* at the end of the simulation, denoted 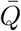, is generally higher when selection on *Q* is strong (Fig. 3B-D, red versus blue curves). Critically, the relationship between 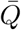 and *N*_*e*_ is not monotonic: 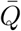 first decreases with *N*_*e*_, but increases when *N*_*e*_ is sufficiently large. Such a relationship indicates different modes of optimization at different *N*_*e*_. When *N*_*e*_ is too small, neither *cis*- nor *trans*-genotypic values can be efficiently optimized, so the starting condition is mostly maintained; when *N*_*e*_ is intermediate, as selection is still not efficient enough to optimize *cis*-loci of individual modification events in the face of mutational pressure and genetic drift, relatively low *Q* evolves to reduce the deleterious effect of gene product modifications globally. When *N*_*e*_ is sufficiently large, selection can have the population approach the global optimum where *Q* is optimal and modification at individual sites are optimized locally via *cis*-substitutions.

The above interpretation predicts that the tipping point where 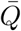 starts to increase with *N*_*e*_ should correspond to a smaller *N*_*e*_ when selection on *Q* is stronger, and that 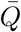 will be lower (given *N*_*e*_) when mutational pressure is strong (i.e., *l* is large) and *cis*-loci are harder to optimize. Both predictions are confirmed by our simulations (Fig. 3B-D). The tipping point occurs at about *N*_*e*_ = 10^2.5^ or *N*_*e*_ = 10^3^ when selection on *Q* is strong (width of fitness function *σ*_*Q*_ = 2; see Methods), but at about *N*_*e*_ = 10^4^ when selection on *Q* is weak (*σ*_*Q*_ = 20). In addition, when *l* is large, 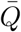 increases less with *N*_*e*_ after the tipping point (Fig. 3B-D).

We also examined how the deleterious modification events are shared across lineages over time. For each modification event, we calculated the fraction of lineages that shared it, and used the median across all 100 modification events to represent the level of conservation given the parameter combination (see Methods). The fraction of lineages sharing the modification generally declined over time, but declined more rapidly when *N*_*e*_ is large and when *l* is small (Fig. 3E-G, Fig. S3). When *N*_*e*_ is relatively small (e.g., *N*_*e*_ *<* 10^3^) and/or *l* is high (e.g., *l* = 10), modifications are shared by a large proportion of, and in some case, all lineages (Fig. 3E-G, Fig. S3). Hence, when selection is too weak given mutational pressure and strength of drift, even deleterious modifications can be readily shared, and sharing of a modification event by divergent lineages may not indicate it is beneficial.

Together, our simulations of *cis-trans* coevolution demonstrate how opposing selection mediated by deleterious gene product modifications and the modification machinery’s additional functions can constrain the optimization of gene product diversity. Such latent functions may explain the maintenance of some known types of gene product diversity. For instance, functional significance of protein recoding by A-to-I RNA editing has been of great interest, but most recoding events do not have any known function and are likely non-adaptive [35, 36]. A-to-I editing in non-coding RNAs transcribed from repetitive elements, however, are involved in preventing autoimmune responses [72–75] and suppressing retrotransposition [76]. In coleoids, where A-to-I editing activity in neural tissues is unusually high [21, 77], A-to-I editing is also enriched in repetitive elements [78], indicating A-to-I editing might perform similar functions in coleoids as well. Another process that is likely explainable by such a model is m6A modification, which is found to be involved in repression of endogenous retroviruses [79] and decay of mis-processed RNA [67] in a mass-action fashion. It should be noted that the above explanation does not indicate there cannot be adaptive modification events in the focal category (e.g., re-coding by A-to-I editing), as it is plausible that adaptive modifications can evolve secondarily with the modification machinery already in place. Similarly, it is compatible with an entrenchment model [43–46] as well, as deleterious substitutions can be permitted and entrenched while the modification machinery is maintained due to its additional function; modifications that “restore” the permitted substitutions can also be considered as a latent function that contributes to the modification process’s maintenance.

Our finding also revealed different optimization strategies at different *N*_*e*_, which is in line with previous findings regarding global and local optimization in the evolution of quality-control mechanisms to reduce fitness cost of expression errors [80–84]. In actual biological systems, the global solution may realize as lowered expression or catalytic efficiency of the *trans*-factor, or an auto-regulatory mechanism where the *trans*-factor modifies its own gene product and trigger negative regulatory effects when its expression is too high [67, 85–87].

### Simulated data recapitulate divergence of A-to-I RNA editing in coleoids

To complement our theoretical results, we asked whether simulation under our model is able to generate a distribution of modification levels that is similar to those observed in empirical studies. To this end, we examined if simulations could recapitulate the distribution of A-to-I RNA editing levels in coleoids. Previous studies reported preponderant A-to-I editing by by the ADAR family of enzymes (adenosine deaminases acting on RNA) in these coleoids’ neural tissues, but the distribution of editing levels at coding sites is strongly skewed, with a vast majority of editing sites having rather low (*<* 1%) editing levels [21, 36, 77]. We simulated evolution of 20,000 editing-type modification events, including 10,000 neutral modifications and 10,000 deleterious modifications along a phylogenetic tree of four coleoid species (Fig. 4A), with some gene-specific parameters (*α, l*, and *C*) sampled from pre-specified distributions. To reproduce a skewed distribution of modification levels like those observed in empirical studies [21, 36, 77], we sampled *C* from an exponential distribution with a moderate mean (i.e., magnitudes higher than *ϵ* but not high enough to produce an editing level above 10%). Editing levels from our simulation showed strong phylogenetic signal (i.e., neighbor-joining tree based on distance in editing levels recapitulates topology of the species tree and relative lengths of branches; Fig. 4B), and has a skewed distribution in each species (exemplified by distribution in octopus shown in Fig. 4C). Similar patterns were seen when neutral (Fig. S5A-B) and deleterious (Fig. S5C-D) editing sites were examined separately, though deleterious editing levels are generally lower. Together, our result demonstrate that observed editing level distributions can be explained by a non-adaptive model where whole-molecule ADAR binding affinity has a skewed distribution across genes.

**Figure 4:**
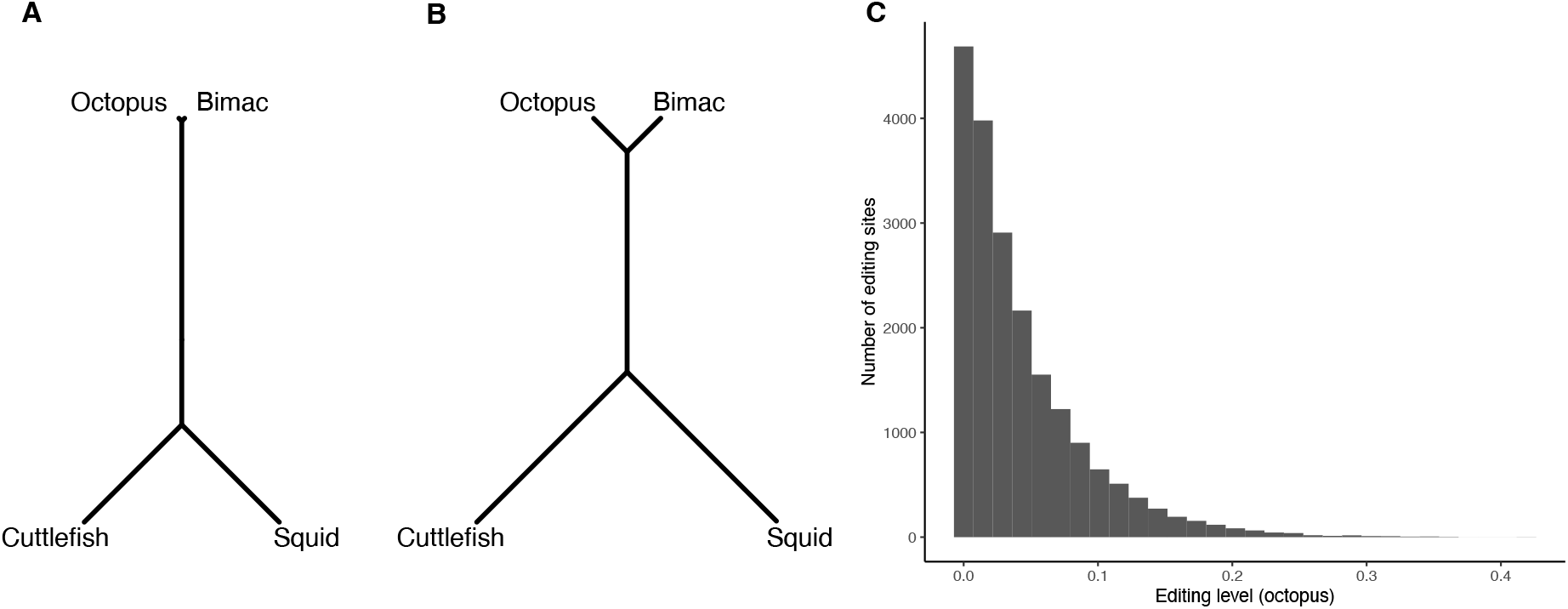
Evolutionary simulations recapitulated patterns of A-to-I RNA editing in four coleoid species, the octopus (*Octopus vulgaris*), the bimac (*O. bimaculoides*), the squid (*Doryteuthis pealeii*), and the cuttlefish (*Sepia oficianalis*). (A) Phylogenetic tree of four coleoid species. (B) Neighbor-joining tree of four coleoid species based on simulated editing levels. An unrooted version is shown in (A) as it is readily comparable to (B). (C) Distribution of editing levels across genes in the octopus.

### Concluding Remarks

In this study, we developed a theoretical model for the evolution of gene product diversity, under which we investigated how the interplay of mutations, genetic drift, and selection on isoform abundances will shape the evolutionary dynamics of gene product diversity. Our analyses of this model revealed that optimization of gene product diversity can be highly constrained by the underlying genetic architecture, the effective population size, expression level of the gene, and pleiotropic effects of the gene product modification machinery— which suggests that a substantial portion of observed gene product diversity is likely to be evolutionarily sub-optimal. Looking forward, it would be informative to conduct more comprehensive empirical analyses across a broader array of taxa; a current impediment to doing so is the lack of statistical phylogenetic tests for comparing the observed distribution of gene product diversity with that expected under the scenarios we studied. While standard statistical approaches for quantitative traits have been shown to be adequate for modeling the evolution of mRNA abundances across a phylogeny [50, 51], enabling direct comparisons between theory and data [53], this is unlikely to hold true for gene product diversity *per se*, owing to its particular genetic and mutational architecture. Our model provides a quantitative framework for developing such statistical tests.

## Methods

### Isoform abundances at equilibrium

Let us consider a scenario where an unmodified isoform (denoted *I*_0_) is converted to a modified isoform (denoted *I*_1_). Their abundances are denoted *P*_0_ and *P*_1_, respectively.

The rate at which *P*_0_ changes through time is given by

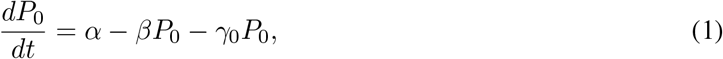

where *α* is the rate at which the unmodified isoform is produced, *β* is the net conversion rate from *I*_0_ to *I*_1_, and *γ*_0_ is the unmodified isoform’s decay rate.

The rate at which *P*_1_ changes through time is given by

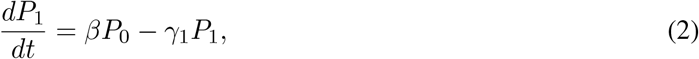

where *γ*_1_ is the modified isoform’s decay rate.

An equilibrium is reached when

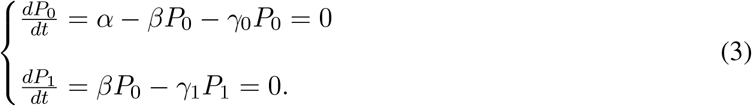

Solving the above system of equations gives

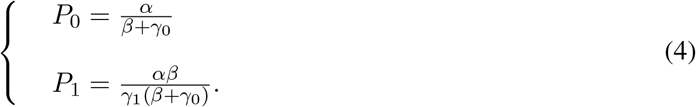

The proportion of the gene product that is modified is

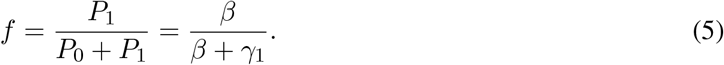

The same model can be extended to more complex cases where more isoforms of the same gene’s product are present. If *n* unique isoforms (*I*_1_, …, *I*_*n*_) can be produced by modifying *I*_0_ and each molecule of *I*_0_ can only be modified into one alternative isoform (i.e., *I*_1_, …, *I*_*n*_ do not convert to each other), the equilibrium is reached when

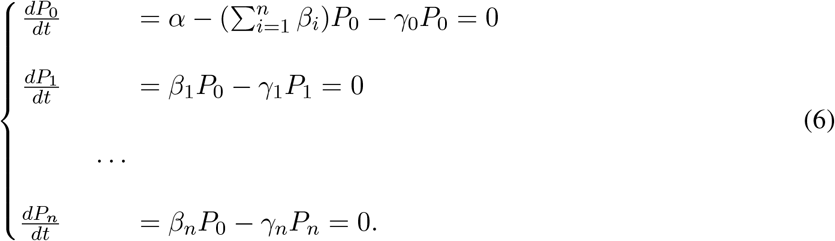

In this case, *β*_1_, …, *β*_*n*_ are net rates at which *I*_0_ is converted to *I*_1_, …, *I*_*n*_, respectively, and *γ*_1_, …, *γ*_*n*_ are decay rates of *I*_1_, …, *I*_*n*_. The above system of equations can be rearranged and written in a matrix (**A***x* = **b**) form:

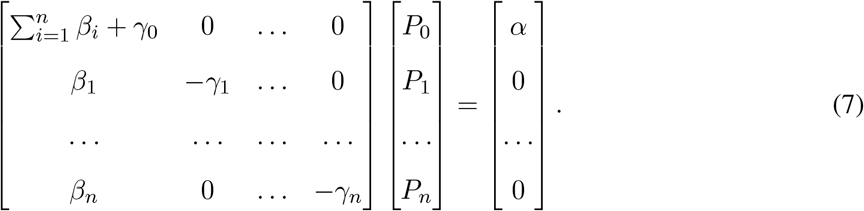

Equilibrium abundances of different isoforms can be obtained by solving the above system of equations.

In this study, we focused on two types of gene product modification processes, editing-type and splicing-type, which are exemplified by RNA editing and splicing, respectively. A variant of the above model is applied to each of the two types. For editing-type modification, we considered a simple case with two isoforms: the unmodified isoform *I*_0_ and the modified isoform *I*_1_. Equilibrium isoform abundances were calculated simply using Eqn. (4). When deriving model predictions, we had *γ*_0_ = 1 and *γ*_1_ = 1, unless stated otherwise. For splicing-type modification, we considered a model with three isoforms: the unmodified isoform *I*_0_ and two modified isoforms, *I*_1_ and *I*_2_. Equilibrium isoform abundances were calculated by solving Eqn. (7) with *n* = 2. When deriving model predictions, we had *γ*_0_ = 0 and *γ*_1_ = 1, unless stated otherwise.

The modeling framework also extends to multi-step modification, where an modified isoform can be further modified into a different one. Let us consider a scenario where a modified isoform *I*_1_ is modified into an different isoform *I*_2_. The equilibrium is reached when

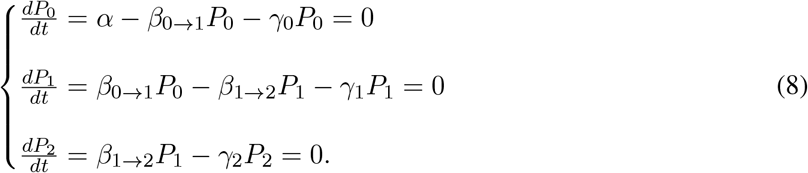

Solving the above system of equations gives

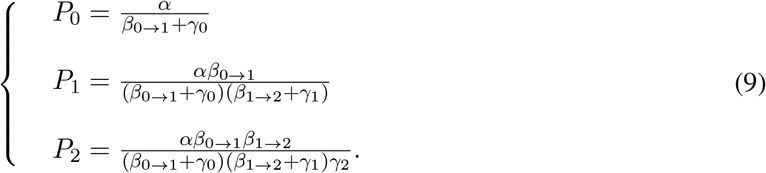

Similarly, if there is a series of *n* modified isoforms where *I*_*i*_ is produced by modifying *I*_*i*−1_:

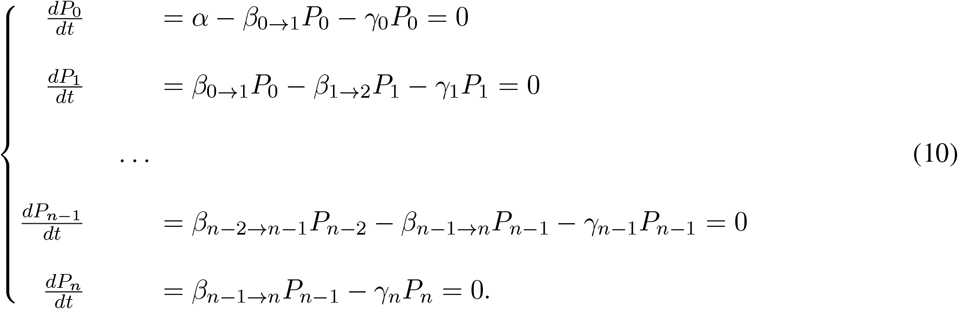

The above system of equations can be rearranged and written in a matrix (**A***x* = **b**) form:

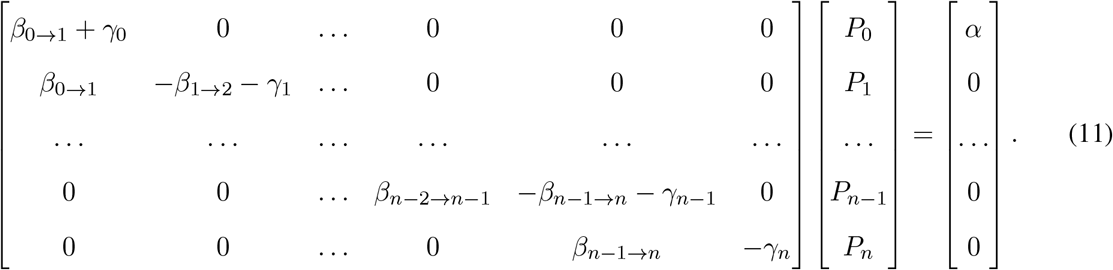

It is worth noting that the above model can be applied when it is the number of modification events within the same RNA or protein molecule but not the exact locations of the modifications that is of interest. In such a case, *n* represents the total number of sites in the RNA or protein molecule that can potentially be modified, and *I*_*i*_ represents isoforms where *i* of the *n* potential sites are modified. If the per-site modification rate is constant regardless of location of the potential modification site or modification states of other sites, such that for each 0 ≤ *i* ≤ *n* − 1 there is *β*_*i*→*i*+1_ = (*n* − *i*)*β*, Eqn. (10) and (11) can be written as

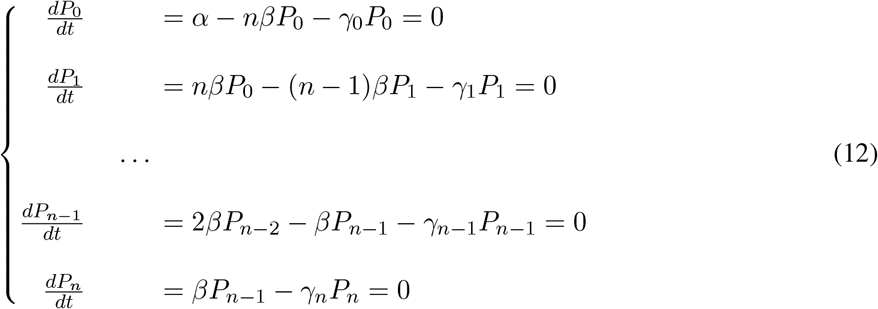

and

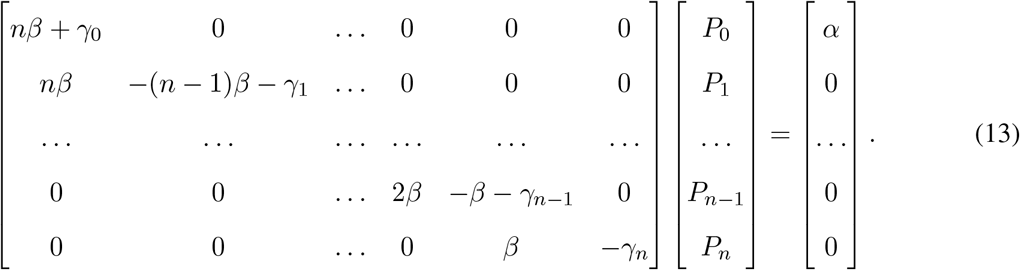

In the most general form of the model where every isoform (e.g., *I*_*i*_) can be converted to another isoform (e.g., *I*_*j*_) at per-molecule rate *β*_*i,j*_ (*β*_*i,j*_ = 0 if *i* = *j*), Eqn. (7) will be written as

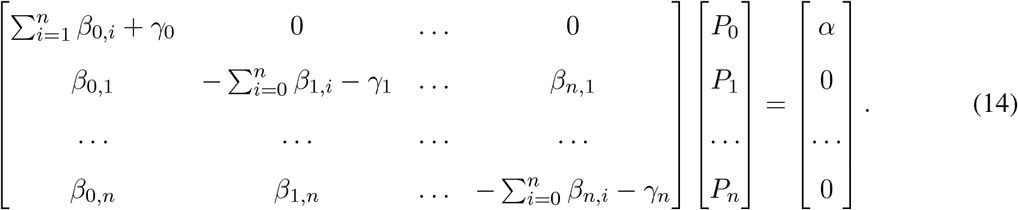

### Genetic architecture of modification rate

For a given modified isoform, the corresponding *β* parameter is determined together by *l cis*-acting loci and a *trans*-genotypic value, *Q*. The *trans*-genotypic value *Q* characterizes the overall activity of the enzyme (or molecular machinery) that carries out the modification process, and is a product of its expression level and per-molecule activity (i.e., the catalysis efficiency of an enzyme, determined by its amino acid sequence and/or conformation). The binding affinity between the enzyme and its substrate is dependent on the *cis*loci, which are sites in regions adjacent to (though not necessarily immediately adjacent to) the site subject to modification.

We assumed that each *cis*-locus can have either an “effector” allele that facilitates binding between the modification enzyme and its substrate (i.e., sequence motif recognized by the enzyme), or a “null” allele that does not facilitate binding. The total effect of the *cis*-loci on *β* is determined by a normalized genotypic value 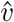, which is calculated as

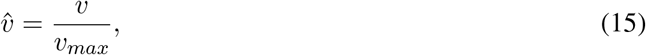

where *v* is the sum of all effector alleles’ effect, and *v*_*max*_ is the greatest possible value of *v* (i.e., achieved when there are no null alleles). In this study, we focused on a simple model where the *cis*-loci’s effect is additive and all *cis*-loci have equal effect, so there *v* is equal to the total number of effector alleles and *v*_*max*_ is equal to the number of *cis*-loci, *l*.

The relationship between *β* and underlying parameters is given by

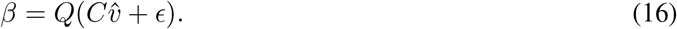

Here, *ϵ* ≥ 0 is the rate of non-specific modification that takes place independent of the *cis*-genotype, and *C >* 0 reflects global structural features of an RNA or protein molecule that affect binding affinity between the enzyme and the substrate.

For splicing-type modification, we assumed that *β*_1_ and *β*_2_ are affected by the same set of *cis*-loci. We also assumed the two alleles that each *cis*-locus could potentially have are both effector alleles. One of them only facilitates the production of *I*_1_, whereas the other only facilitates the production of *I*_2_; under this model, the same genotype’s effects on *I*_1_ and *I*_2_ are inversely correlated. For convenience, we defined the normalized *cis*-genotypic value based on the genotype’s effect on *β*_1_. The *β* parameters are thus given by

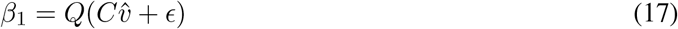

and

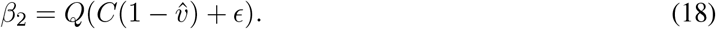

For editing-type modification, we had *C* = 1, *Q* = 1, and *epsilon* = 0 when deriving model predictions, unless specified otherwise. For splicing-type, we had *C* = 1, *Q* = 100, and *epsilon* = 0, unless specified otherwise.

The mutational spectrum of each *cis*-locus is characterized by two per-locus mutation rates: *μ*_01_, the rate of mutations from the null allele to the effector allele, and *μ*_10_, the rate of mutations in the opposite direction. The difference between *μ*_01_ and *μ*_10_ reflects difference in the two allele’s corresponding sequence spaces (i.e., number of nucleotide states, assuming each locus represents a single site) and/or rate of different types of nucleotide changes (e.g., transition/transversion bias or AT-bias). In the case of splicing-type modification, *μ*_01_ and *μ*_10_ are simply replaced by mutation rates between two effector alleles. For simplicity, we assumed that all *cis*-loci have the same mutational spectrum in this study. In this study, we had the total mutation rate per locus *μ*_01_ + *μ*_10_ = 2 *×* 10^−9^, unless stated otherwise.

### Selection on isoform abundance

We first considered a scenario where each isoform contributes to fitness independently, in which case the fitness is given by

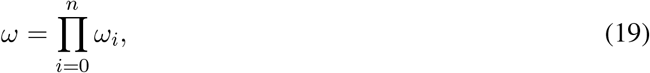

where *ω*_*i*_ is fitness with respect to *P*_*i*_.

We considered two scenarios where an isoform’s abundance is subject to selection: a scenario where the isoform is functional and a scenario where it is not functional but deleterious. Selection on the abundance of a functional isoform *I*_*i*_ is characterized by a Gaussian fitness function:

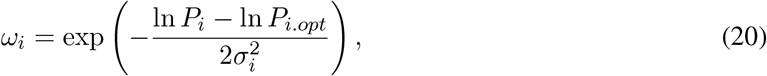

where 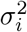, the fitness function’s width and characterizes the strength of selection.

If *I*_*i*_ is deleterious, fitness with respect to its abundance *P*_*i*_ is given by

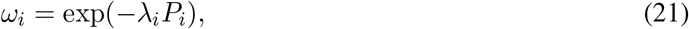

where *λ*_*i*_ *>* 0 is a parameter characterizing the strength of selection. When *λ*_*i*_ = 0, there is *ω*_*i*_ = 1, which corresponds to the case that *P*_*i*_ is not under selection. In this study, we had *σ* = 10 for every functional isoform and *λ* = 10^−3^ for every deleterious isoform, unless specified otherwise.

For editing-type modifications, we mainly focused on a scenario where the modification is deleterious– i.e., *I*_0_ is functional while *I*_1_ is toxic. Fitness component with respect to *P*_0_ is calculated by Eqn. 20), whereas fitness with respect to *P*_1_ is calculated by Eqn. 21). For splicing-type modifications, fitness is determined only by *P*_1_ and *P*_2_, not *P*_0_. One of the modified isoforms, *I*_1_, is the functional, and its abundance *P*_1_ is under stabilizing selection; fitness component with respect to *P*_1_ is thus computed using Eqn. 20). The other modified isoform, *I*_2_, in contrast, is not functional but toxic, and the corresponding fitness component is computed using Eqn. 21). With *I*_2_ representing mis-processed isoform(s), we also assumed that *γ*_2_ is greater than *γ*_1_ to recapitulate quality-control mechanisms that act to eliminate mis-processed isoforms [62– 64]; specifically, we examined scenarios of *γ*_1_ = 1 while *γ*_2_ is equal to 20, 50, or 100.

### Distribution of *cis*-genotypic value

When the number of *cis*-loci underlying a modification event is reasonably small, the evolution of genotypic value *v* (and thus 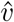) can be approximated by a sequential-fixation (strong-selection-weak-mutation) model [88]. Then, assuming that other parameters that affect modification are constant, the evolution of *v* (and 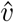) can be modeled as a Markov process with a constant transition matrix. A time step in this Markov process can be a generation or any arbitrary time interval as long as the the probability that more than one mutations arise in the population is very low (i.e., 2*N*_*e*_*μ <* 0.01) such that the sequential-fixation model is an appropriate approximation [29]. Using this approach, the distribution of the *cis*-genotypic value *v* given the starting state after a given amount of time can be derived.

Let us consider a simple scenario where the effector allele at every *cis*-locus has an effect size of 1 (i.e., *v* is equal to the number of effector alleles and *v*_*max*_ = *l*). In a diploid population, the probability that *v* becomes *v* + 1 via a substitution in a time step given the present genotypic value *v* is

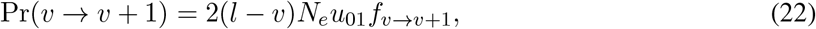

where *N*_*e*_ is the effective population size and *f*_*v*→*v*+1_ is the fixation probability given ancestral and mutant phenotypes.

Similarly, the probability of becoming *v* − 1 via a substitution is

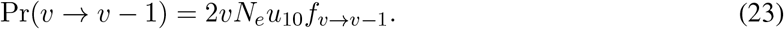

The probability that *v* does not change is simply

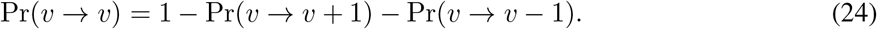

The fixation probability is obtained using Kimura’s method [89]:

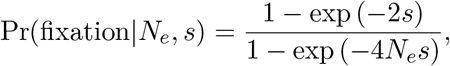

where 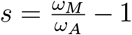 is the coefficient of selection (*ω*_*M*_ and *ω*_*A*_ represent mutant and ancestral fitness, respectively).

Given the probability distribution of *v* at a time *t*, **v**_*t*_, the distribution at *t* + 1 is

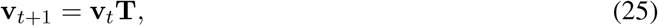

where **v**_*t*_ and **v**_*t*+1_ are row vectors of length *l* + 1, with each element represents the probability of a possible value of *v*. The transition matrix **T** is a *l* + 1 *× l* + 1 matrix where **T**[*i* + 1, *j* + 1] = Pr(*i* → *j*). The probability Pr(*i* → *j*) is calculated following Eqn. 22 and 23 if 0 ≤ *i* ≤ *l*, 0 ≤ *j* ≤ *l*, and |*i* − *j*| ≤ 1; otherwise, Pr(*i* → *j*) = 0. In this study, we used **v**_0_**T**^1e8^ to represent an equilibrium distribution. For editing-type modification, we had the first element of **v**_0_ equal to 1 (i.e., starting from the genotype that has the least effect on modification), whereas for splicing-type modification, we had the last element of **v**_0_ equal to 1 (i.e., starting from the genotype that maximizes the production of *I*_1_ and minimized the production of *I*_2_).

If different *cis*-loci have different effect sizes, there will be up to 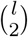 possible values of *v*. In the extreme case where all loci have different effect sizes and the mutation rate varies depending both on the locus and the ancestral allele, the transition probability from a given genotype to a given neighbor genotype (i.e., differ from the ancestral genotype at only one site) is simply the product of the mutation rate (given the locus and the ancestral allele) and the fixation probability. In this study, we focused mostly on the simple scenario where all the *cis*-loci have equal effect size and mutation rates, although the modeling framework can be readily extended to more general cases.

### Simulating *cis-trans* coevolution

To investigate coevolutionary dynamics between the *cis*-loci and the *trans*-genotypic value *Q* when many genes or sites are subject to modification, we conducted simulations of evolution where *cis*-loci and *Q* are both affected by mutations.

Each lineage we simulated was divided into a number of time steps, with the number of time steps proportional to the branch length. If the only loci that could undergo evolutionary changes in a time interval are the *cis*-loci, the probability distribution of a given modification event’s *cis*-genotypic value *v* at the end of the time interval is simply

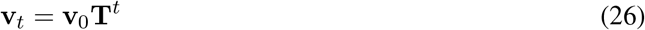

where *v*_0_ is the starting distribution and *t* is the number of time steps the time interval consists of. If the simulation starts from a pre-designated value of *v*, the corresponding element of **v**_0_ will be 1 while other elements are equal to 0.

Before simulating evolution for a lineage, we first determined *m*, the total number of mutations that affect *Q* to occur during evolution by sampling *m* from a Poisson distribution with mean equal to 2*N*_*e*_*U*_*Q*_*L*, where *L* is the branch length (i.e., number of time steps) and *U*_*Q*_ is the rate of mutations that affect *Q*. Then we randomly picked *m* time steps, at each of which a mutation affecting *Q* would occur. If *m > L* (which has very low probability given parameter values considered, and did not happen in our simulations), this value of *m* will not be used for simulations. The effect of each mutation on ln *Q* was sampled from a normal distribution *N* (0, S_*Q*_). Change in the distribution *v* during the interval between two mutations that affect *Q* obtained using Eqn. 26, with *t* being the number of time steps between two mutations. Before examining the fitness effect of a mutation that affects *Q*, a value of *v* was first sampled from its distribution, which, together with the mutation’s effect on *Q*, will determine the fixation probability. If the mutation is fixed, the transition matrix will be re-calculated with the mutant *Q*, and the mutant *Q* will be the new *Q* to begin with when the next mutation is examined. When products of multiple genes are subject to modification, fitness effect of each mutation affecting *Q* is determined collectively by its effect on all modification events; when such a mutation is fixed, all gene’s transition matrices will be altered. For simplicity, we assumed that different modification events’ *cis*-loci are not shared and evolve independently (i.e., no linkage between *cis*-mutations affecting different modification events).

We considered a scenario where the modification machinery has both beneficial and detrimental effects on fitness at the same time. Under this model, there are a set of genes subject to deleterious editing-type modifications (i.e., the unmodified isoform is functional and the modified isoform is deleterious). At the same time, *Q* contributes to a fitness component *ω*_*Q*_ that is independent of these modification events. In our simulations, *Q* was under stabilizing selection, and *ω*_*Q*_ is given by

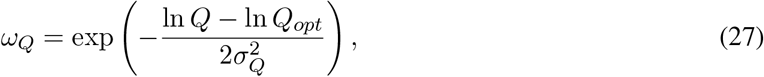

where *Q*_*opt*_ is the optimal value of *Q* and *σ*_*Q*_ is the fitness function’s width. In this case, if there are *n* genes subject to modification, the overall fitness is given by

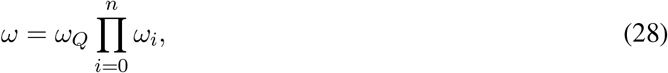

where *ω*_*i*_ is fitness with respect to the *i*-th gene’s isoform abundances.

Values of *N*_*e*_ used in the simulations include 10^2^, 10^2.5^, 10^3^, 10^3.5^, 10^4^, 10^4.5^, and 10^5^. In each simulation, we considered 100 genes that are subject to deleterious modifications. For simplicity, we had all modification events have equal *l*, and considered scenarios of *l* = 2, *l* = 5, and *l* = 10, where the initial value of *v* was 1, 2, and 5, respectively. Regarding selection on *Q*, we considered two scenarios: a scenario of strong selection (*σ*_*Q*_ = 2) and a scenario of relatively weak selection (*σ*_*Q*_ = 20). In all simulations, we had *Q*_*opt*_ = 2, *U*_*Q*_ = 10^−8^ and S_*Q*_ = 0.1. We also had *α* = 1, *γ*_0_ = 1, *γ*_1_ = 1, *C* = 1, *σ* = 10, *λ* = 10^−3^, and *ϵ* = 10^−3^ for all genes in all simulations. Starting value of *Q* was equal to its optimum for all simulations. After the simulations, we quantified the degree to which the modifications are shared (i.e., conserved) among lineages. For each gene, we calculated the the fraction of lineages where *P*_1_ *>* 0.005. The median of all genes is then used to represent how likely a modification event is shared given the evolutionary parameters (*l, N*_*e*_, and strength of selection). We examined how this value varied depending on divergence time by performing the simulation with different times of duration, including 2 *×* 10^7^, 4 *×* 10^7^, 6 *×* 10^7^, 8 *×* 10^7^, and 10^8^ time steps. For each combination of parameter values, we simulated 50 independent lineages.

The above procedure can also be used to simulate the coevolution of the *cis*-loci and other parameters, such as *α, C* or *ϵ*, in which case mutations affecting *Q* in the above procedure will be replaced by mutations affecting the parameter of interest.

### Simulation along the coleoid tree

We simulated evolution of editing levels at 20, 000 editing sites along a phylogenetic tree of four coleoid species: the common octopus (*Octopus vulgaris*), the bimac (*O. bimaculoides*), the squid (*Doryteuthis pealeii*), and the cuttlefish (*Sepia oficianalis*). The coleoids have high A-to-I RNA editing activity in their neural tissues, whereas extant non-coleoid cephalopods and non-cephalopod mollusks do not [21, 77]. Branch lengths of the phylogenetic tree are based on divergence times described in ref. [21], with mid point the reported range used for our simulations. Divergence time of the octopus and the bimac, which are very closely related, was set to be 5 million years. We assumed each time step in the simulation corresponds to a year, so the number of time steps a branch corresponds to is equal to branch length in terms of years. We started the simulation from the most recent common ancestor of four coleoids, and the value of *v* of each editing site at this ancestral node was sampled randomly from the corresponding genotypic space. We assumed that *Q* is under strong stabilizing selection mediated by functions independent of the focal editing events such that *Q* remained constant in the simulation. We had *Q* = 1 for this simulation. The distribution of *v* at the end of each branch was obtained using Eqn. (26) with time of evolution equal to branch length; a value of *v* was then sampled from the distribution to represent the state at the end of this branch and the starting state of its descendent branches (if any). Some gene-specific parameters were sampled from pre-specified distributions. The rate at which *I*_0_ is expressed, *α*, was sampled from a log-normal distribution; that is, ln *α* was sampled from *N* (0, 1). The number of *cis*-loci, *l*, was sampled uniformly from (0, 1, …, 10). The *C* parameter was sampled from a exponential distribution with mean equal to 0.1. All genes had *γ*_0_ = 1, *γ*_1_ = 1, *σ* = 10, *λ* = 10^−3^, and *ϵ* = 10^−4^. Because *ϵ >* 0, all editing levels were positive. Thus, after the simulation, we log-transformed all editing levels and computed Euclidean distances between each pair of species using log-transformed editing levels (ln (*f*)). We then built a neighbor-joining (NJ) tree based on these distances using the *nj* function of R package *ape*, and asked this NJ tree recapitulate the phylogenetic relationship of the four coleoid species; specifically, we examined whether 1) the two *Octopus* species fall in one clade while the squid and the cuttlefish fall in another, and 2) whether distance between the two octopuses is closer than that between the squid and the cuttlefish.

## Code and data availability

Code and data files are available at https://github.com/phylo-lab-usc/gene_product_diversity

## Acknowledgements

We thank Mark Kim, and members of the Pennell, Edge, and Mooney labs for their thoughtful comments on parts of this study. We acknowledge support from the Natural Sciences and Engineering Research Council of Canada (FN 492860, to AFP), the Jean D’Alembert Foundation (France 2030 program ANR-11-IDEX-0003, to AFP), and the National Institute of General Medical Sciences (R35GM151348, to MP).

## Tables

**Table 1:** Definitions and notations of parameters.

| Parameter | Definition |
| --- | --- |
| $I_i$ | The $i$-th modified isoform; $I_0$ represents the unmodified isoform. |
| $P_i$ | Abundance of $I_i$. |
| $\alpha$ | Rate at which $P_0$ is produced. |
| $\beta_i$ | Per-molecule net rate at which the $I_0$ is converted to the $I_i$. |
| $\gamma_i$ | Decay rate of $I_i$. |
| $f$ | Modification level; $f = \frac{P_1}{P_1+P_0}$ for editing-type and $f = \frac{P_2}{P_1+P_2}$ for splicing-type. |
| $l$ | Number of *cis*-loci affecting $\beta$. |
| $v$ | *cis*-genotypic value characterizing the combined effect of the *cis*-genotype on $\beta$. |
| $v_{max}$ | Value of $v$ when every locus has an effector allele. |
| $\hat{v}$ | Normalized *cis*-genotypic value, $\hat{v} = \frac{v}{v_{max}}$. |
| $Q$ | *trans*-genotypic value underlying $\beta$. |
| $C$ | Parameter characterizing gene-level feature that affect *cis*-loci's effect size on $\beta$. |
| $\mu_{01}$ | Mutation rate from null allele to effector allele per *cis*-loci. |
| $\mu_{10}$ | Mutation rate from null allele to effector allele per *cis*-loci. |
| $\omega$ | Overall fitness. |
| $\omega_i$ | Fitness with respect to $P_i$. |
| $\sigma_i$ | Width of the fitness function when $P_i$ is under stabilizing selection. |
| $\lambda_i$ | Parameter characterizing speed at which $\omega_i$ declines with $P_i$ when $I_i$ is toxic. |
| $s$ | Coefficient of selection of a mutation. |
| $N_e$ | Effective population size. |
| $\Pr(i \rightarrow j)$ | Probability that $v$ changes from $i$ to $j$ via a substitution in a time step. |
| $\mathbf{T}$ | Transition matrix for the genotypic value $v$. |
| $\mathbf{v}_t$ | Probability distribution of $v$ at a given time $t$. |
| $v_t$ | Value of $v$ at a given time $t$; $v_0$ is the starting value. |
| $U_Q$ | Rate of mutations affecting $Q$. |
| $S_Q$ | Standard deviation of mutation's effect on $\ln Q$. |
| $\sigma_Q$ | Width of the fitness function of $Q$. |

## Supplementary materials

**Figure S1:**
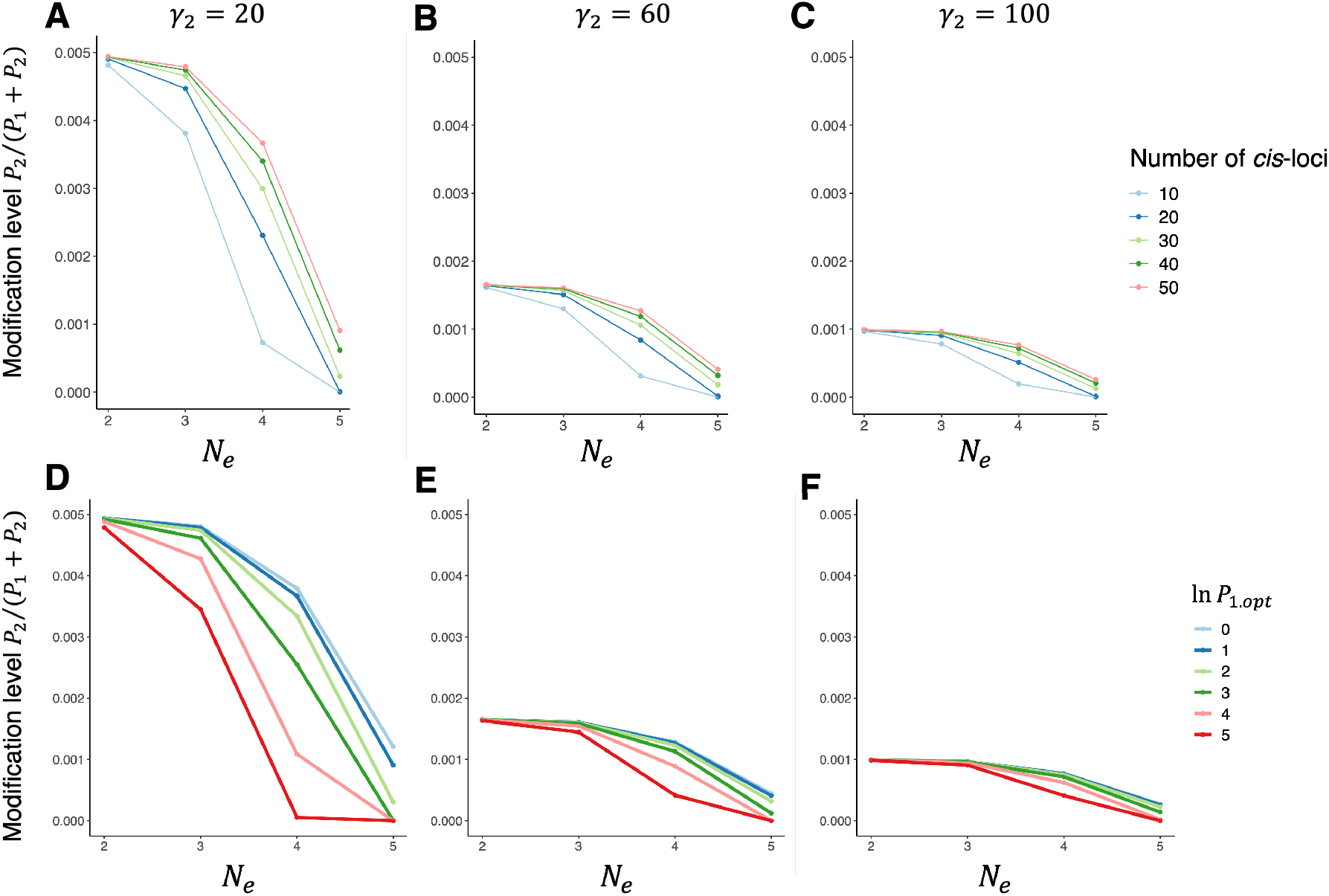
Scaling between mean modification level of splicing-type modification and effective population size *N*_*e*_ (shown in log10 scale). (A-C) Response of mean modification level to *N*_*e*_ under different combinations of *cis*-loci number (*l*) and decay rates of the dysfunctional isoform (*γ*_2_), with with optimal expression level *P*_1.*opt*_ = exp (1) (ln *P*_1.*opt*_ = 1). (D-F) Response of mean modification level to *N*_*e*_ under different 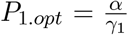 and *γ*_2_, with *l* = 50. All results are derived with initial *cis*-genotypic value *v*_0_ = *l* (i.e., maximizing *β*_1_ and minimizing *β*_2_), time of evolution *T* = 10^8^ time steps, *μ*_01_ = *μ*_10_ = 10^−8^, *Q* = 100, *γ*_0_= 0,*γ*_1_ = 1, and 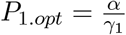 .

**Figure S2:**
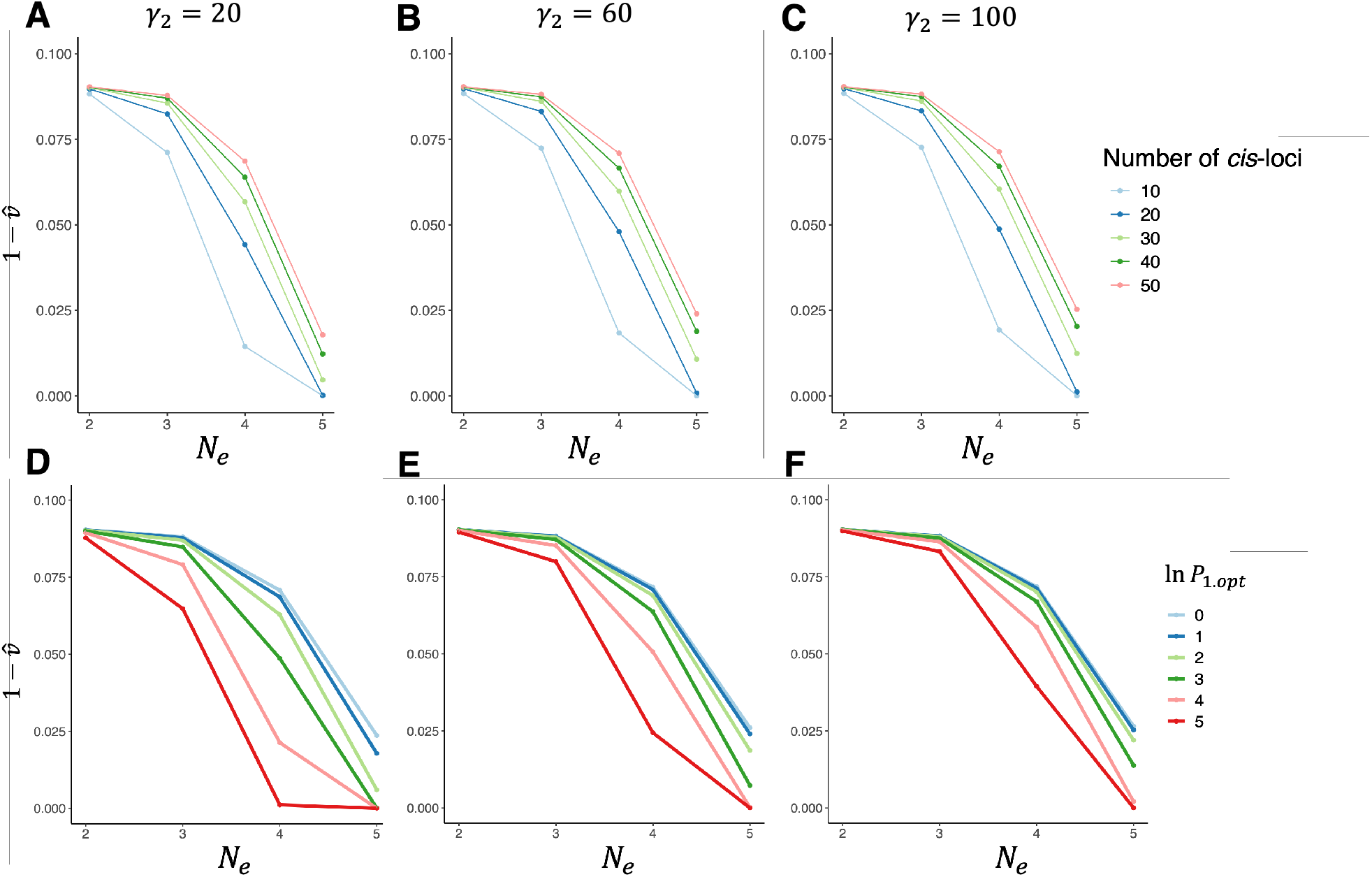
Scaling between normalized mean *cis*-genotypic value of splicing-type modification and *N*_*e*_ (shown in log10 scale). Represented by the Y-axes is 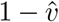, which reflects the degree to which *cis*-genotype favors production of the dysfunction and toxic isoform *I*_2_. (A-C) Response of 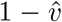 to *N*_*e*_ under different combinations of *l* and *γ*_2_, with optimal expression level *P*_1.*opt*_ = exp (1) (ln *P*_1.*opt*_ = 1). (D-F) Response of 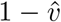 to *N*_*e*_ under different *P*_1.*opt*_ and *γ*_2_, with *l* = 50. All results are derived with initial *cis*-genotypic value *v*_0_ = *l*, time of evolution *T* = 10^8^ time steps, and *μ*_01_ = *μ*_10_ = 10^−8^, *Q* = 100, *γ*_0_ = 0, *γ*_1_ = 1, and 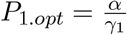.

**Figure S3:**
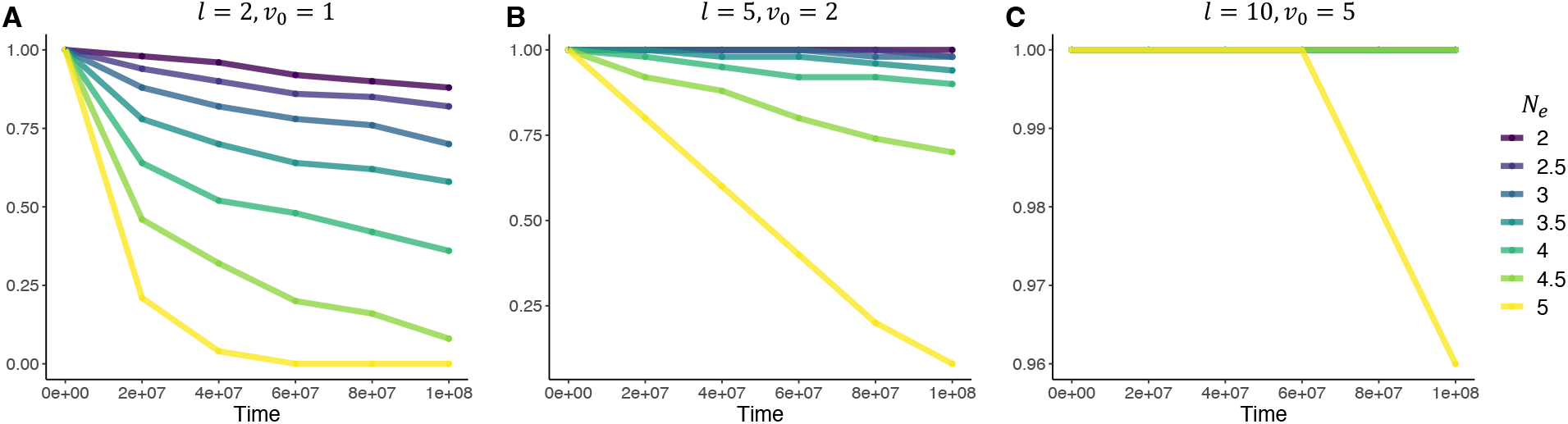
Sharing of modification events over time. Y-axes represent among-gene median of proportion of lineages (species) that share a modification event when selection on *Q* is weak (*σ*_*Q*_ = 20). When two curves in the same panel completely overlap, the one with the largest corresponding *N*_*e*_ is shown.

**Figure S4:**
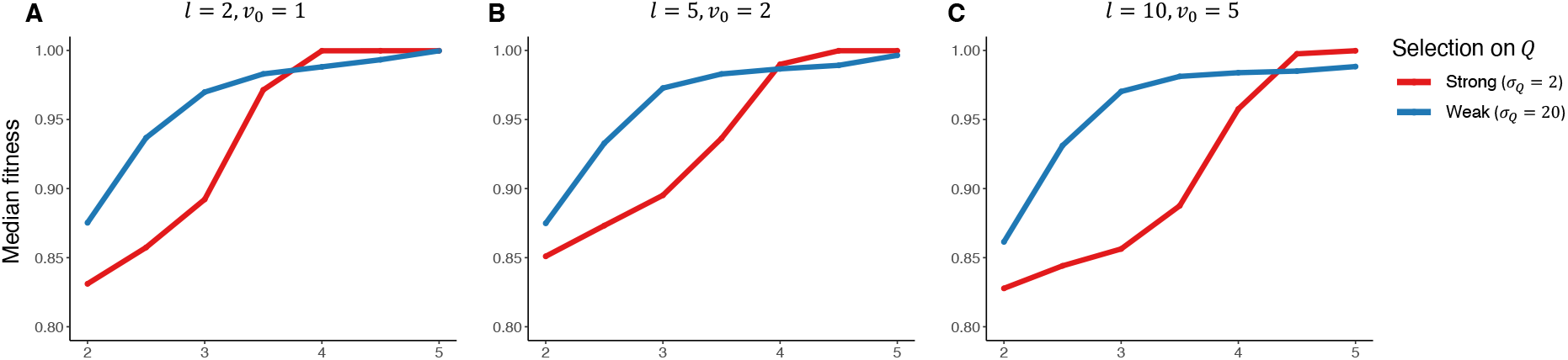
Response of median fitness across lineages (species) to *N*_*e*_ when the modification process is under opposing selection forces. (A) *l* = 2, *v*_0_ = 1. (B) *l* = 5, *v*_0_ = 2. (C) *l* = 10, *v*_0_ = 5. Maximum fitness (fitness value at the global optimum) is equal to 1.

**Figure S5:**
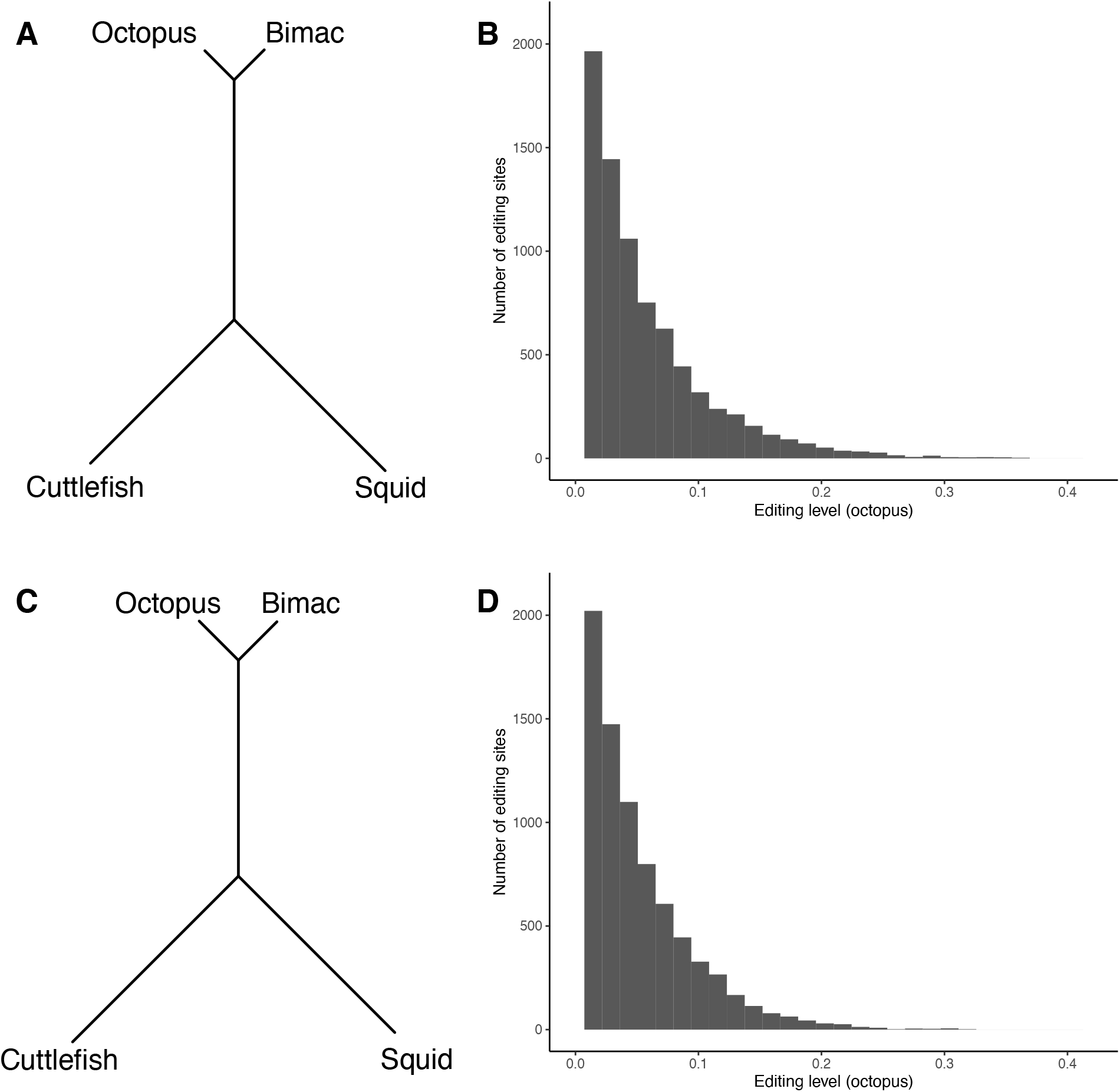
(A) Neighbor-joining tree of four coleoid species based on simulated neutral editing levels. (B) Distribution of neutral editing levels in the octopus. (C) Neighbor-joining tree of four coleoid species based on simulated deleterious editing levels. (D) Distribution of deleterious editing levels in the octopus.

